# Bat Red Blood Cells express Nucleic Acid Sensing Receptors and bind RNA and DNA

**DOI:** 10.1101/2022.01.13.476238

**Authors:** LK Metthew Lam, Jane Dobkin, Kaitlyn A. Eckart, Ian Gereg, Andrew DiSalvo, Amber Nolder, Eman Anis, Julie C. Ellis, Greg Turner, Nilam S. Mangalmurti

## Abstract

Red blood cells (RBCs) demonstrate immunomodulatory capabilities through the expression of nucleic acid sensors. However, little is known about bat RBCs, and no studies have examined the immune function of bat erythrocytes. Here we show that bat RBCs express the nucleic acid-sensing Toll-like receptors TLR7 and TLR9 and bind the nucleic acid ligands, single-stranded RNA, and CpG DNA. Collectively, these data suggest that, like human RBCs, bat erythrocytes possess immune function and may be reservoirs for nucleic acids. These findings provide unique insight into bat immunity and may uncover potential mechanisms by which virulent pathogens in humans are concealed in bats.

## Introduction

A fundamental role of the immune system is nucleic acid-sensing, which is essential for pathogen detection and coordination of host defense.^1,2^ Toll-like receptors (TLRs) are critical for nucleic acid recognition and include TLR9, which detects DNA containing unmethylated CpG motifs present in bacterial, viral, and mitochondrial DNA (mt-DNA), and TLR7, which recognizes single-stranded RNA (ssRNA) present in viruses and host RNA. Most studies on TLR function have focused on their roles in classical immune cells, although some enucleated cells such as platelets also express TLRs.^3,4^ Because mammalian red blood cells (RBCs) are devoid of organelles, the prevailing assumption has been that they lack immune function. Contrary to this dogma, a DNA-sensing role for RBCs through the expression of TLR9 has recently been discovered.^5,6^ Whether bat RBCs bind nucleic acids or express TLRs is unknown.

Members of the order Chiroptera (bats) are exceptional in their ability to harbor deadly viruses without developing disease.^7^ Although substantial evidence supports a bat origin for numerous viruses, how bats tolerate such viruses and other pathogens remains unknown.^8^ Bats are hosts to diverse intraerythrocytic haemosporidian parasites and co-evolved with these parasites.^9,10^ Additionally, as an adaption to flight, the number of erythrocytes per microliter of blood is two to five-fold higher than in humans, increasing the total red cell surface area.^11^ Despite these observations, little is known regarding bat erythrocytes and their interaction with the immune system. Because we identified TLRs on mammalian RBCs, we asked if bat RBCs similarly express TLRs and bind nucleic acids.

## Results

### Bat RBCs Display Diverse Morphologies

Big brown bats (*Eptesicus fuscus*) are a hibernating North American micro-bat species that commonly use anthropogenic structures for hibernation and maternal colonies, even to some degree in winter, and are frequently submitted to wildlife rehabilitation facilities throughout the year, often presenting with injuries. We obtained terminal blood samples from wild-caught big brown bats admitted to Pennsylvania wildlife rehabilitators that were determined to be non-releasable due to their conditions (Table 1). All bats were consistently in a euthermic state during their time at the facility. Samples were collected post-euthanasia following transfer to the Pennsylvania Game Commission. We first prepared peripheral blood smears to examine RBC morphology under basal conditions (Fig.1a). Consistent with previous reports, we noted crenated and poikilocytic cells (Fig.1b).^11^ Unlike human RBCs, where crenated RBCs were primarily observed following DNA binding and during acute inflammatory states, the number of abnormal RBCs was equivalent to normal-appearing RBCs in bats that spent sufficient time in rehabilitation (Fig. 1c).^12^ We also noted a substantial amount of heterogeneity in the appearance of crenated cells. Two of the bats demonstrated significant hemolysis and the presence of intra-erythrocytic inclusions, marked by erythrocyte ghosts and Howell-Jolly bodies, respectively (Fig. 1d, Supplemental Fig. 1).

**Table 1.** Big brown bat (*Eptesicus fuscus*) subjects.

| **Bat** | **Sex** | **Weight (g)\*** | **Days in Rehabilitation** | **Clinical Presentation** |
| --- | --- | --- | --- | --- |
| **A** | M | 9.8 | 91 | Injured wing |
| **B** | F | 11.2 | 70 | Grounded due to unknown cause |
| **C** | F | 16.5 | 27 | Grounded and captured at remodeling work site |
| **D** | M | 11.7 | 33 | Multiple membrane holes and fracture to right wing, small distal fracture to left wing, possible frost bite |
| **E** | F | 13.5 | 98 | No visible injuries |
| **F** | M | 12.7 | 6 | No visible injuries |
| **G** | M | 9.4 | 4 | Shredded membrane and open fracture of left wing |
| **H** | F | 12.0 | 8 | Shredded membrane and exposed bone of right wing |
| **I** | M | 11.3 | 9 | Alopecia along left shoulder, unspecified injury to left wing, degloving injury to one toe of left foot |
| **J** | M | 20.9 | 193 | Injury and bruising to right leg, cuts along left leg, hip, and back |
| **K** | M | 16.4 | >240 | No visible injuries but unexplained significant weight loss despite healthy appetite |

\* Weight was measured at time of euthanasia.

**Figure 1.**
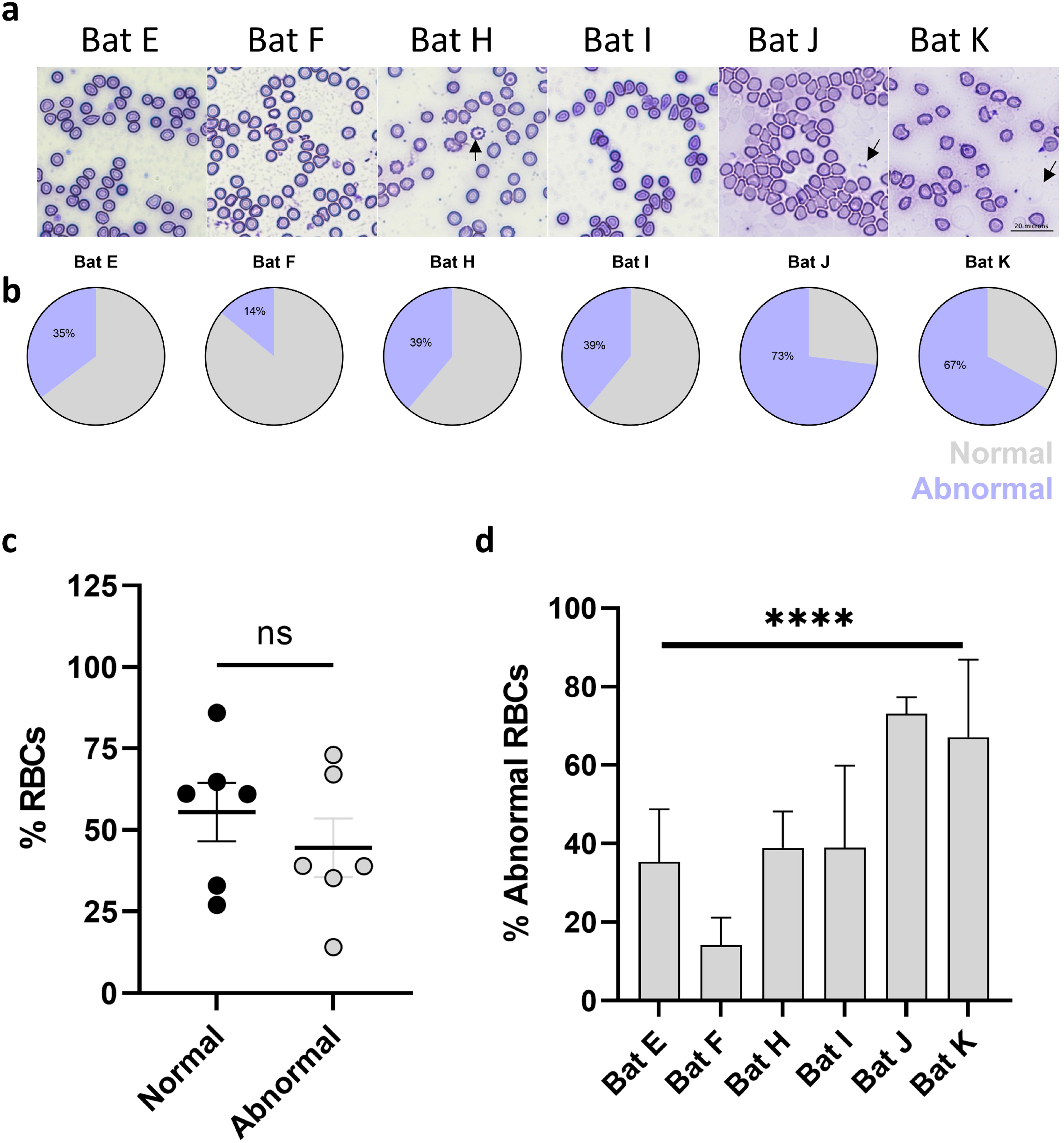
Morphological Characteristics of Bat RBCs. a. Peripheral smears of RBCs from six bats. b. percentage of normal and abnormal RBCs in individual bats. c. similar numbers of normally shaped and abnormally shaped RBCs are observed in bats. d. percentage of abnormal RBCs in bats, ****P<0.0001. Arrows denote erythrocyte ghosts.

### Nucleic Acids Bind to Bat RBCs

Next, we measured the ability of bat RBCs to bind RNA and DNA. Fluorescently labeled GU-rich single-stranded RNA (ssRNA, RNA-40) and CpG oligodeoxynucleotides (ODNs) were incubated with bat RBCs before washing and analyzing by flow cytometry. Consistent with human RBCs, bat RBCs bound both RNA and CpG ODNs in a dose-dependent fashion (Figure 2a-c). Because microbial DNA binds to human RBCs, we assayed purified bat RBCs for the presence of RNA or DNA by performing qPCR for 18S rRNA and the 16S rDNA gene.^12^ When compared with RBCs from healthy human donors, bat RBCs contained significantly less bacterial DNA but similar amounts of 18S RNA(Supplemental Fig. 2a and b). Taken together, these data would suggest that bat RBCs exhibit differential nucleic acid binding when compared with human RBCs. Since a bat origin of SARS-CoV-2 has been proposed, we asked whether bat RBCs would bind RNA from SARS-CoV-2. Although only one animal was available for studies, we observed robust binding of SARS-CoV-2 RNA to bat RBCs (Supplemental Fig. 2c and d). Interestingly, previous studies have demonstrated that inoculation of big brown bats (*Eptesicus fuscus)* with SARS-CoV-2 both orally and nasally does not lead to infection, viral shedding, or antibody generation.^13^ The most likely explanation for these findings is the low expression of the receptor for SARS-CoV-2 entry, ACE-2. However, another plausible explanation would be the rapid clearance of the virus by circulating RBCs. Given the extremely thin alveolar-capillary barrier present in Chiropterans, it is possible that inhaled microbes may transit into the circulation, although further studies are needed to support this theory.^13,14^

**Figure 2.**
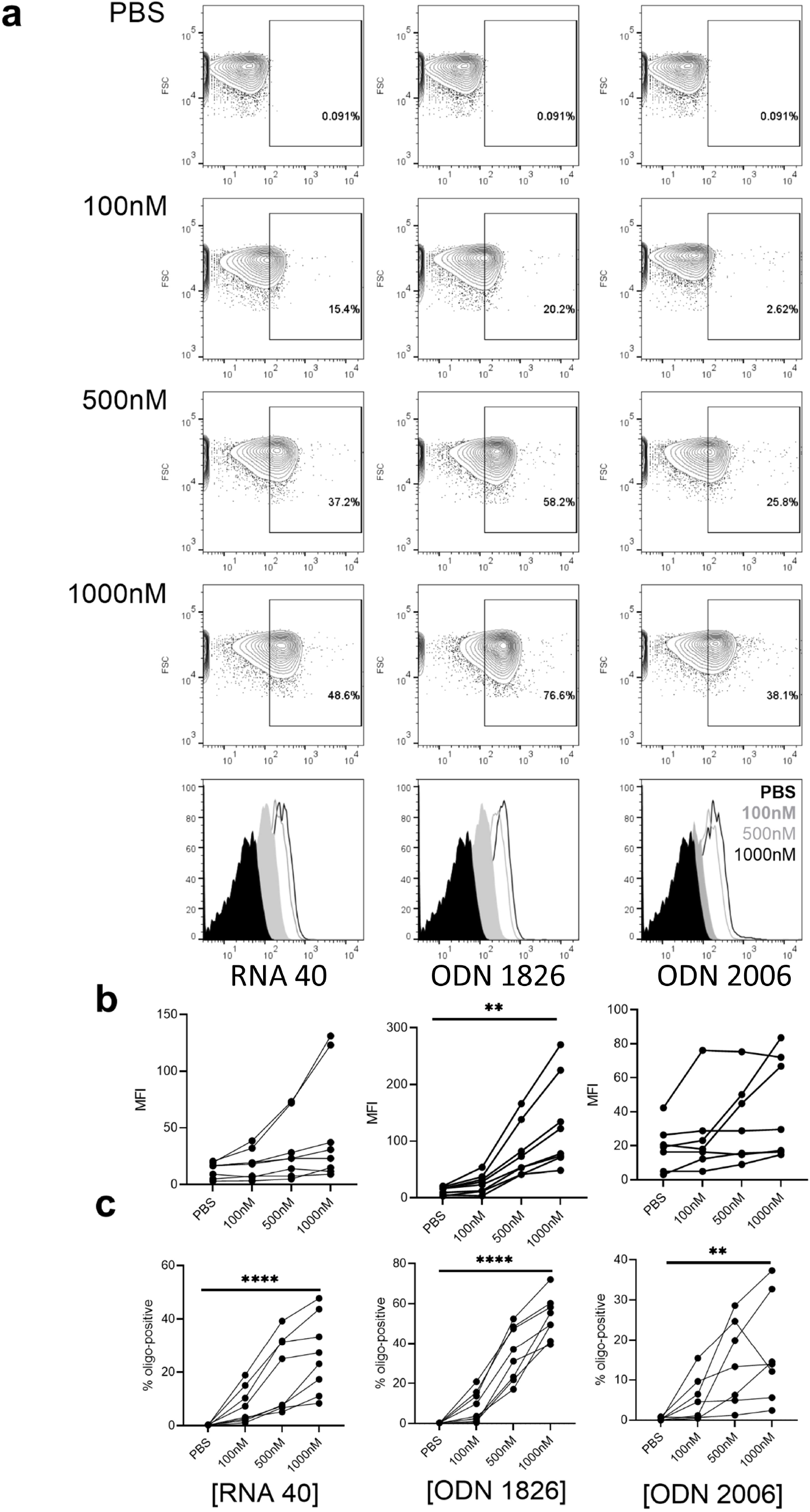
Bat RBCs bind RNA and DNA. a. dose dependent binding of ssRNA (RNA 40), CpG ODN 1826 or 2006, contour plots and histogram are shown, one representative bat. b. Summary statistics, geometric Mean Fluorescence Intensity (MFI) and percent positive cells (c), n=8 bats. **P<0.01, ****P<0.0001.

### Bat RBCs Express TLR7 and TLR9

We next asked if CpG-binding would alter bat RBCs. Consistent with previous studies, CpG acquisition by RBCs led to marked alteration of the bat RBCs, Supplemental Fig. 3.^5,15^ Concurring with earlier observations, naive untreated bat RBCs also displayed a range of morphologies, Supplemental Figure 3a. Nucleic acid sensing in bats is a topic of intense investigation and the toll-like receptors (TLRs) have been extensively characterized in bats.^16,17^ We, therefore, asked if bat RBCs expressed TLRs. Flow cytometry of intact, unpermeablized bat RBCs did not identify TLRs on the RBC surface, Supplemental Fig. 4. However, staining of permeabilized cells did reveal the presence of TLR9 and TLR7, although TLR3 was not detected, Figure 3a-b. We confirmed these findings with immunofluorescence, Figure 3c. However, only four of the five bats tested consistently demonstrated positive TLR7 and TLR9 staining, suggesting heterogeneity in erythroid TLR expression amongst mammals. Notably, the bats with the highest TLR7 staining were also the bats nearest to their injury before arrival to the rehabilitation facility (Table 1).

**Figure 3.**
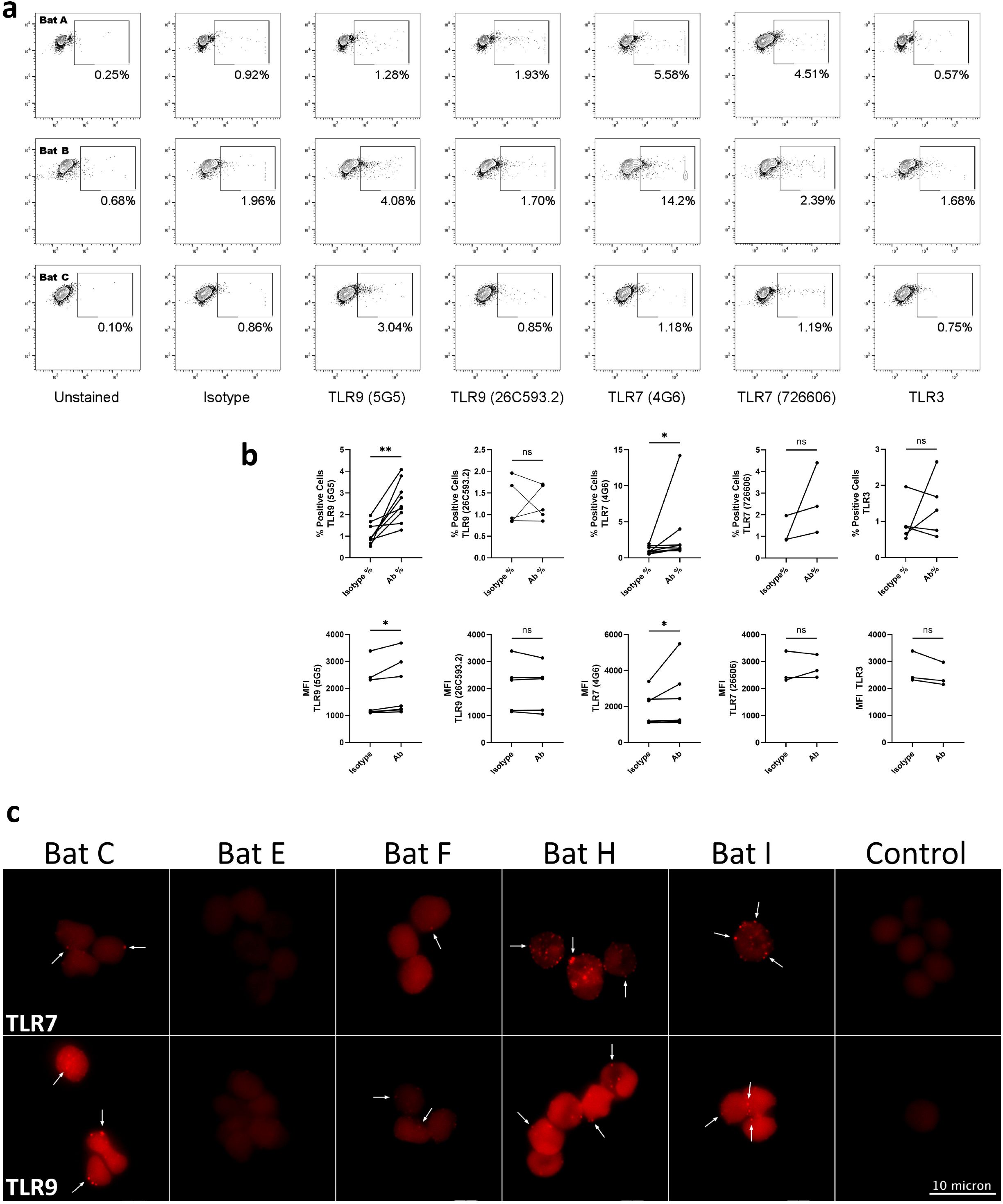
TLR expression on Bat RBCs. a. Intracellular staining for TLR 9,7 and 3 from three representative bats. b. Summary statistics, percent positive cells and MFI for TLRs, n=5, *P<0.05, **P<0.01. c. Immunofluorescent staining for TLR7 and TLR9. Arrows denote positive signal (bright red punctae).

## Discussion

Collectively our data demonstrate the presence of nucleic acid-sensing TLRs on bat RBCs and nucleic acid-binding capabilities of bat RBCs. Furthermore, bat RBCs contain RNA, suggesting that these RBCs may serve as reservoirs for RNA. As a result of the preponderance of bat-borne viral pathogens with pandemic potential, substantial interest in bat immunity has emerged. Recently an immune function of RBCs through nucleic acid-sensing and delivery to classical immune cells has been identified.^9^ Given the large number of circulating RBCs present in Chiropteran species and the uniquely rich lipid composition of bat RBCs, we speculate that lipid-rich bat RBCs function to shelter RNA from the immune system, permitting host tolerance.^18^ Interestingly, in rehabilitated bats, RBCs exhibited a substantial number of abnormal cells and features, including excessive hemolysis, poikilocytes, Howell-Jolly bodies, and crenated cells, reminiscent of those found in hemolytic anemias, acute infection, and RBC membrane disorders, including pyropoikilocytosis and elliptocytosis. Additionally bat TLRs display mutations in ligand binding sites that may alter their ability to bind nucleic acids, yet whether these adaptations affect erythrocyte TLRs remains an open question.^16,17^ Because bats undergo dramatic temperature changes from hypothermia during torpor to elevated temperature during flight, it is possible that the unique RBC features are not just a consequence of acclimatization to pathogens but also an adaption to rapid dynamic temperature changes, as has been observed in some congenital human red cell disorders such as pyropoikilocytosis.^19^ The extreme metabolic stress of flight has also driven red cell adaptations including increased hematocrit and smaller RBCs allowing for greater oxygen transport and thus may also have influenced other RBC changes.^14^ Although the driver of these unique RBC alterations in Chiroptera remain unclear, our findings of nucleic acid binding by bat RBCs suggests an immune function for these RBCs.

The discovery of an immune role for enucleate, mature RBCs prompts the question of what evolutionary pressure led to such adaptations. Mounting evidence suggests that human non-gas exchanging RBC adaptations arose in response to the dominant selective pressure of malaria-causing parasites of the genus *Plasmodium*, with chemokine binding through DARC (Duffy Antigen Receptor for Chemokines) and hemoglobin S (sickle cell) being the two most prominent examples. Mammals in the order Chiroptera co-exist with *Plasmodium* and other haemosporidians, and some have postulated a Chiropteran origin to primate malaria.^10^ Future studies of mammalian erythrocyte immune function, including nucleic acid binding and complement regulation in bats and other mammalian hosts, will be necessary to elucidate the complete picture of red cell immune function, potentially providing insight into mechanisms by which pathogens lethal to humans are tolerated in other mammals.

## Methods

### Animals and ethic statements

Animal studies were conducted using residual blood obtained from bats in rehabilitation and provided by the Pennsylvania Game Commission. All studies were done in accordance with the Institutional Animal Care and Use Committee. Studies using human specimens were approved by the Institutional Review Board. Healthy volunteers provided informed consent prior to study participation.

### Bat blood collection

Bats were anesthetized with isoflourane and then euthanized via cervical dislocation. Blood was collected either via cardiac puncture or from major vessels after decapitation and placed in EDTA microtainers. Samples were centrifuged for 15 min at 3,500 rpm (max RCF = 1,534g) and then transferred to the University of Pennsylvania for further analysis.

### Human RBC Isolation

Healthy volunteers provided informed consent. Whole blood was collected in EDTA tubes prior to centrifugation at 3000g for 10 min. The plasma and buffy coat were aspirated. Red blood cells were purified from the remaining packed red cell fraction using magnetic-assisted cell sorting (MACS) as previously described.^12^

### Blood processing and RBC isolation

Whole blood was centrifuged at room temperature for 5min at 800g. Plasma and buffy coat were isolated and saved. The remaining packed red blood cells (RBCs) were resuspended in 1mL PBS for washing. Then, the cells were centrifuged, and the supernatant was aspirated along with a small amount of the packed RBCs on the top. To determine the yield of RBCs, serially diluted packed RBCs were counted on a hemacytometer. Immediately following this, the RBCs were either used for functional assays, fixed, or frozen in aliquots. To freeze RBCs, 10^7^ RBCs were added to DNA lo-bind tubes and pelleted by centrifugation at 10,000 rpm for 5min. The supernatant was removed and the cell pellets were store at −80°C. Residual packed RBCs were stored in Adsol (Fenwal) at 3:1 volume ratio at 4°C.

### RBC smears and staining

To generate smears for RBCs, 5μL of PBS was added to a clean Superfrost Plus microscope slide (Fisherbrand) and 2μL of washed packed RBCs were mixed with the PBS and spread along the slide. The slides were air-dried and stored at room temperature until usage. Air-dried smears were stained using DiffQuik.

### Nucleic acid binding assay

250,000 RBCs in 100μL sterile PBS were transferred to round-bottom polystyrene tubes (Falcon, 352052). Afterwards, oligonucleotides diluted in sterile PBS were added to the cells in a volume of 5μL. Sequences for the oligonucleotides are listed in Methods Table 1. The tubes were placed on a rack, sealed with parafilm, covered in aluminum foil, and rocked on a nutator at 37°C for 2hr. To stop the binding reactions, the cells were washed in 1mL PBS and pelleted at 800g for 5min with slow deceleration. A total of two washes were performed. The cells were resuspended in 300μl PBS and analyzed by flow cytometry (BD Fortessa). Immediately prior to data acquisition, the cells were resuspended by pipetting up and down repeatedly.

**Methods Table 1.** Oligonucleotides used in binding assays.

| Name | Sequence | Source |
| --- | --- | --- |
| RNA40-ATTO647 | rG*rC*rC*rC*rG*rU*rC*rU*rG*rU*rU*rG*rU*rG*rU*rG*rA<br>*rC*rU*rC-ATTO647NN | IDT |
| ATTO647-ODN2006 | ATTO647NN-<br>T*C*G*T*C*G*T*T*T*T*G*T*C*G*T*T*T*G*T*C*G*<br>T*T | IDT |
| FITC-ODN2006 | AlexaFluor488N-<br>T*C*G*T*C*G*T*T*T*T*G*T*C*G*T*T*T*G*T*C*G*<br>T*T | IDT |
| APC-ODN1826 | AlexaFluor647-<br>T*C*C*A*T*G*A*C*G*T*T*C*C*T*G*A*C*G*T*T | IDT |
| FITC-ODN1826 | T*C*C*A*T*G*A*C*G*T*T*C*C*T*G*A*C*G*T*T-FITC | Invivogen |

\* denotes phosphothiorate bond; r denotes ribonucleotide
IDT, Integrated DNA technology, Inc.

### Viral RNA binding assays

SARS-CoV-2 RNA was a kind gift provided by Dr. Kellie Jurado, University of Pennsylvania. 10^7^ RBCs in 100μL sterile PBS were transferred to 2mL DNA/RNA lo-bind tube (Eppendorf, 86-924) and mixed with indicated amount of viral RNA diluted in sterile PBS. The mix was incubated at 37°C for 2hr on a nutator. At 1hr after incubation, tubes were rotated to ensure cells remain resuspended. After the incubation, cells were washed with 1mL sterile PBS three times and frozen at −80°C prior to RNA extraction.

### Surface antibody staining

The cells were resuspended in FACS buffer (PBS + 2% FBS) to 10^7^ cells/mL. For each staining reaction, 10^6^ RBC were pelleted in round bottom polystyrene tubes and stained in 100μL of an antibody mix diluted in FACS buffer. Information for the antibodies used are in Methods Table 2. The cells were rocked at room temperature for 1hr in the dark. The cells were washed with 1mL PBS twice as described above and analyzed by flow cytometry.

**Methods Table 2.** Antibodies against TLR used.

| **Target** | **Clone** | **Usage:**<br>**dilution** | **Source** | **RBC condition** | **Notes** |
| --- | --- | --- | --- | --- | --- |
| TLR3 | 40C1285.6 | FC: 5µg | Novus Biological | Fresh intact cells<br>Fixed and permeabilized | Very few positive cells noted. |
| TLR7 | 4G6 | IFA: 1:200<br>FC: 1:200 | Novus Biological | Fixed and permeabilized | Positive staining both IFA and FC. |
| TLR7 | 726606 | IFA: 1:200<br>FC: 5µg | R&D Systems | Fixed and permeabilized | No detection by IFA. Few positive cells FC. |
| TLR9 | 5G5 | FC: 5µg | Abcam | Fresh cell surface stain or fixed and permeabilized | Positive staining. |
| TLR9 | ab37154 | IFA: 1:200 | Abcam | Fixed and permeabilized | Positive staining. |
| TLR9 | 26C593.2 | FC: 5µg | Novus Biological/Abcam | Fixed and permeabilized | Very few positive cells noted. |

FC: flow cytometry; IFA: immunofluorescent assay

### RBC fixation

RBCs were resuspended to 10^7^ cells/mL in PBS, and an equal volume of 0.1% glutaraldehyde diluted in PBS was added to the cells for a final concentration of 0.05%. The cells were fixed at room temperature in the dark for 10min. Fixation was stopped by adding equal volume of FACS buffer to the cells. The cells were pelleted at 800g for 5min and washed a total of three times. For long-term storage, fixed cells were resuspended in FACS supplemented with 0.05% sodium azide and stored at 4°C.

### Intracellular staining with flow cytometry

10_6_ RBCs were used in each staining reaction. Fixed RBCs were permeabilized in 0.1% TritonX-100 diluted in PBS for 15min at room temperature. The cells were washed and pelleted in FACS buffer, blocked in PBST (PBS + 0.05% tween20) supplemented with 1% BSA for 1hr, and then pelleted and resuspended in antibody diluted in PBST. Information for the antibodies used is listed in Methods Table 2. For flow cytometry staining, cells were incubated with either antibody or isotype controls diluted in FACS buffer for 1hr in the dark at room temperature under gentle shaking conditions. Subsequently, the cells were washed with FACS buffer twice and stained with a secondary antibody diluted in PBST for 45min at room temperature. The cells were then washed in FACS buffer three times prior to analysis.

### Immunofluorescent staining

RBCs were fixed, permeabilized, and blocked as described above. The primary antibody was diluted in PBST with 1% BSA and incubation was performed overnight at 4°C. Subsequently, cells were washed in PBST three times and resuspended in a secondary antibody diluted in PBST. After 1hr of staining at room temperature, the cells were washed in PBST three times. Following the final wash, the buffer was aspirated, and the residual volume of the RBCs was further reduced by pipetting. The remaining cells were resuspended in no more than 10μL PBS. To mount cells on a slide, no more than 5μL of Fluoromount G was added to a microscope slide and 5μL RBCs were mixed in it, and samples were sealed with a coverslip.

### Microscopy and quantification

All micrographs were taken on Nikon A2 microscope. Manual counting quantification was performed by at least two personnel, one of which is blinded.

### DNA extraction

10^7^ Frozen RBCs were thawed on ice. DNA was extracted using DNeasy blood/tissue kit (Qiagen) according to the manufacturer’s protocol. DNA was eluted in 152μL elution buffer at the final step.

### RNA extraction and reverse transcription

1×10^7^ Frozen RBCs were thawed on ice. RNA was extracted using RNeasy Plus Kit (Qiagen) according to the manufacturer’s protocol. During the initial step, RBCs were lysed in 600μL lysis buffer and during the final step, RNA was eluted in 30μL water. 8μL of freshly isolated RNA was reverse transcribed to cDNA via the Superscript First Strand Synthesis system (ThermoFisher) following the manufacturer’s protocol.

### Imagestream

Adsol-stored RBCs were washed in PBS three times. 7×10^5^ RBCs were then incubated with the specified oligonucleotides at 100nM or 1000nM at a final volume of 200uL for 2hr at 37°C. Cells were washed in PBS twice and fixed in 0.05% glutaraldehyde for 10min. Prior to Imagestream (Amnis) analysis, the cells were washed in FACS buffer twice. Analysis was performed using IDEAS software. The automated feature finder was used to discriminate between RBC populations of different morphology. The “aspect ratio” and DNA binding differentiated populations that displayed distinct RBC morphology.

### qPCR

Nucleic acids were quantified via the QuantStudio7 Flex System. For 18S rRNA, cDNA were amplified in PowerUp SYBR Green master mix with the primers (18S rRNA F: AACCCGTTGAACCCCATT and 18S rRNA-R: CCATCCAATCGGTAGTAGCG). For bacterial 16S rDNA, DNA was amplified in Taqman Fast Universal Mastermix with the primers (BSF8: AGA GTT GAT CCT GGC TCA G and BSR357: CTG CTG CCT YCC GTA) and probe (/56-FAM/TA A+CA +CAT G+CA +AGT +GGA /3BHQ_1/ where + denotes locked nucleic acid)

### Statistical Analysis

Statistical analyses were performed with GraphPad Prism 9. Data normality of was determined by Wilk-Shapiro test. Differences among groups were determined with t-test or one-way ANOVA wherever appropriate.

## Acknowledgements

We would like to thank Robyn Grabowski and Rebekah Jones at Centre Wildlife Care and Stephanie Stronsick at Pennsylvania Bat Rescue for their care of the animals and providing whole blood from the bats. We thank Dr. Kellie Jurado for providing SARS-CoV-2 RNA and Dr. Christopher A. Hunter for editorial assistance.

## References

1 Diebold, S. S., Kaisho, T., Hemmi, H., Akira, S. & Reis e Sousa, C. Innate antiviral responses by means of TLR7-mediated recognition of single-stranded RNA. Science 303, 1529–1531, doi:10.1126/science.1093616 (2004).

2 Hemmi, H. et al. A Toll-like receptor recognizes bacterial DNA. Nature 408, 740–745, doi:10.1038/35047123 (2000).

3 Aslam, R. et al. Platelet Toll-like receptor expression modulates lipopolysaccharide-induced thrombocytopenia and tumor necrosis factor-alpha production in vivo. Blood 107, 637–641, doi:10.1182/blood-2005-06-2202 (2006).

4 Semple, J. W., Italiano, J. E. & Freedman, J. Platelets and the immune continuum. Nature Reviews Immunology 11, 264–274, doi:10.1038/nri2956 (2011).

5 Hotz, M. J. et al. Red Blood Cells Homeostatically Bind Mitochondrial DNA through TLR9 to Maintain Quiescence and to Prevent Lung Injury. Am J Respir Crit Care Med 197, 470–480, doi:10.1164/rccm.201706-1161OC (2018).

6 Lam, L. M. et al. Erythrocytes Reveal Complement Activation in Patients with COVID-19. medRxiv, 2020.2005.2020.20104398, doi:10.1101/2020.05.20.20104398 (2020).

7 Hayman, D. T. S. Bat tolerance to viral infections. Nature Microbiology 4, 728–729, doi:10.1038/s41564-019-0430-9 (2019).

8 Schountz, T., Baker, M. L., Butler, J. & Munster, V. Immunological Control of Viral Infections in Bats and the Emergence of Viruses Highly Pathogenic to Humans. Frontiers in immunology 8, 1098, doi:10.3389/fimmu.2017.01098 (2017).

9 Perkins, S. L. & Schaer, J. A Modern Menagerie of Mammalian Malaria. Trends in parasitology 32, 772–782, doi:10.1016/j.pt.2016.06.001 (2016).

10 Schaer, J. et al. High diversity of West African bat malaria parasites and a tight link with rodent Plasmodium taxa. Proc Natl Acad Sci U S A 110, 17415–17419, doi:10.1073/pnas.1311016110 (2013).

11 Schinnerl, M., Aydinonat, D., Schwarzenberger, F. & Voigt, C. C. Hematological survey of common neotropical bat species from Costa Rica. Journal of zoo and wildlife medicine : official publication of the American Association of Zoo Veterinarians 42, 382–391, doi:10.1638/2010-0060.1 (2011).

12 Lam, L. K. M. et al. DNA binding to TLR9 expressed by red blood cells promotes innate immune activation and anemia. Science Translational Medicine 13, eabj1008, doi:doi:10.1126/scitranslmed.abj1008 (2021).

13 Hall, J. S. et al. Experimental challenge of a North American bat species, big brown bat (Eptesicus fuscus), with SARS-CoV-2. Transboundary and emerging diseases, doi:10.1111/tbed.13949 (2020).

14 Maina, J. N. What it takes to fly: the structural and functional respiratory refinements in birds and bats. The Journal of experimental biology 203, 3045–3064 (2000).

15 Lam, L. K. M. et al. Erythrocytes identify complement activation in patients with COVID-19. Am J Physiol Lung Cell Mol Physiol 321, L485–l489, doi:10.1152/ajplung.00231.2021 (2021).

16 Escalera-Zamudio, M. et al. The evolution of bat nucleic acid-sensing Toll-like receptors. Molecular ecology 24, 5899–5909, doi:10.1111/mec.13431 (2015).

17 Jiang, H. et al. Selective evolution of Toll-like receptors 3, 7, 8, and 9 in bats. Immunogenetics 69, 271–285, doi:10.1007/s00251-016-0966-2 (2017).

18 Gillett, M. P. & Wilson, R. B. Unusual lipid composition of erythrocytes from the insectivorous bat Molossus molossus. Comparative biochemistry and physiology. A, Comparative physiology 80, 149–150, doi:10.1016/0300-9629(85)90531-6 (1985).

19 Da Costa, L., Galimand, J., Fenneteau, O. & Mohandas, N. Hereditary spherocytosis, elliptocytosis, and other red cell membrane disorders. Blood Rev 27, 167–178, doi:10.1016/j.blre.2013.04.003 (2013).

