## Supplemental Figures for "Bat Red Blood Cells express Nucleic Acid Sensing Receptors and bind RNA and DNA"

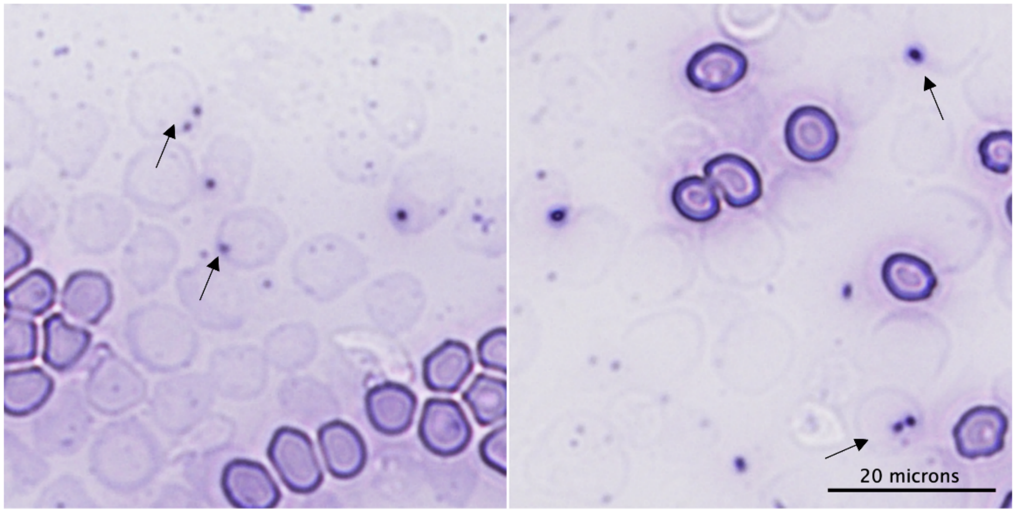


**Supplemental Figure 1. Howell- Jolly Bodies and erythrocyte ghosts are present in bat smears.** Peripheral smears of bat RBCs reveal a significant number of Howell-Jolly Bodies (arrows) and RBC ghosts, indicative of incomplete DNA expulsion and hemolysis.

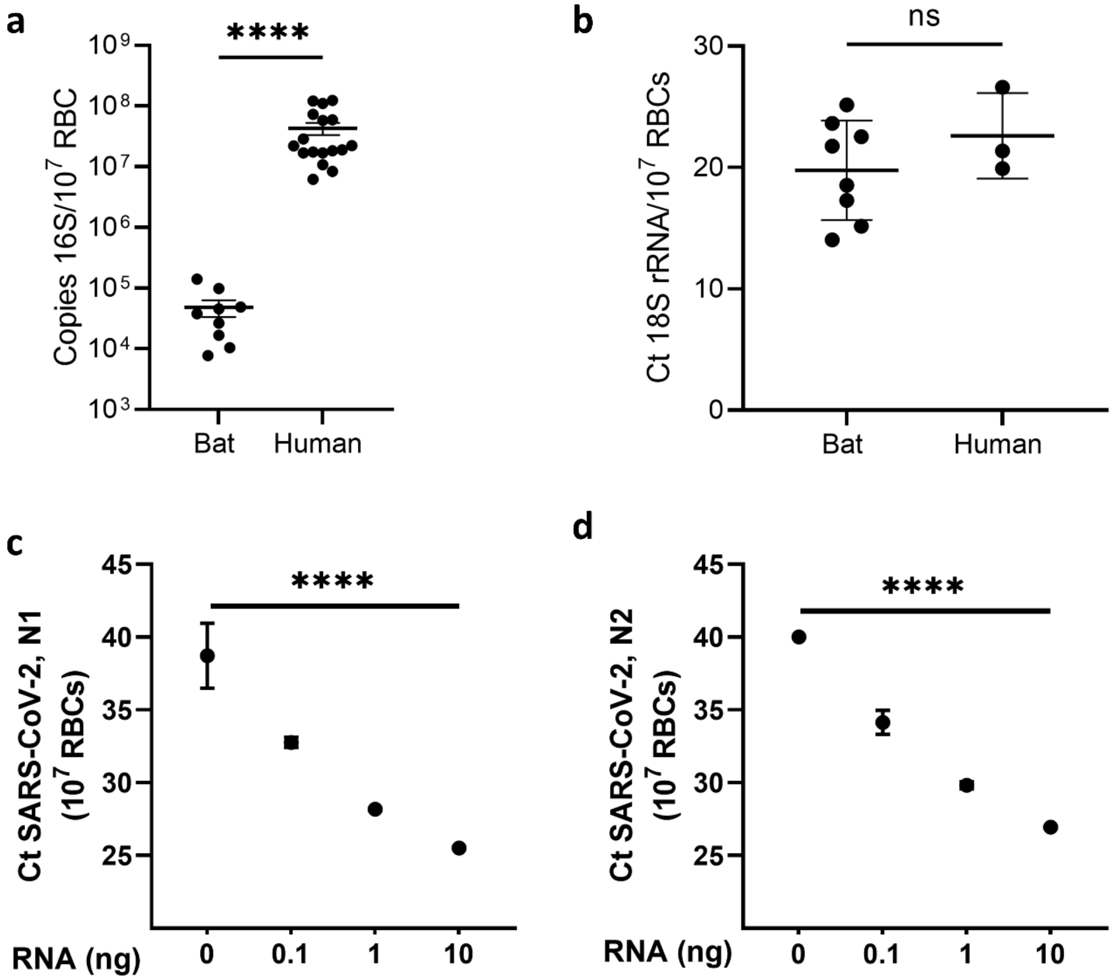


**Supplemental Figure 2. Bat RBCs contain nucleic acids and bind pathogen derived RNA.** a. 16S rRNA gene copies associated with RBCs derived from healthy human donors and bats. b. 18S rRNA content of bat and human RBCs. c and d. qPCR for SARS-CoV-2 on RBCs incubated with varying concentrations of SARS-CoV-2 RNA, PCR for the nucleocapsid protein 1 (N1) and nucleocapsid protein 2 (N2). ****$P<0.0001$, n=1 bat, technical replicates shown for each dose of RNA used.

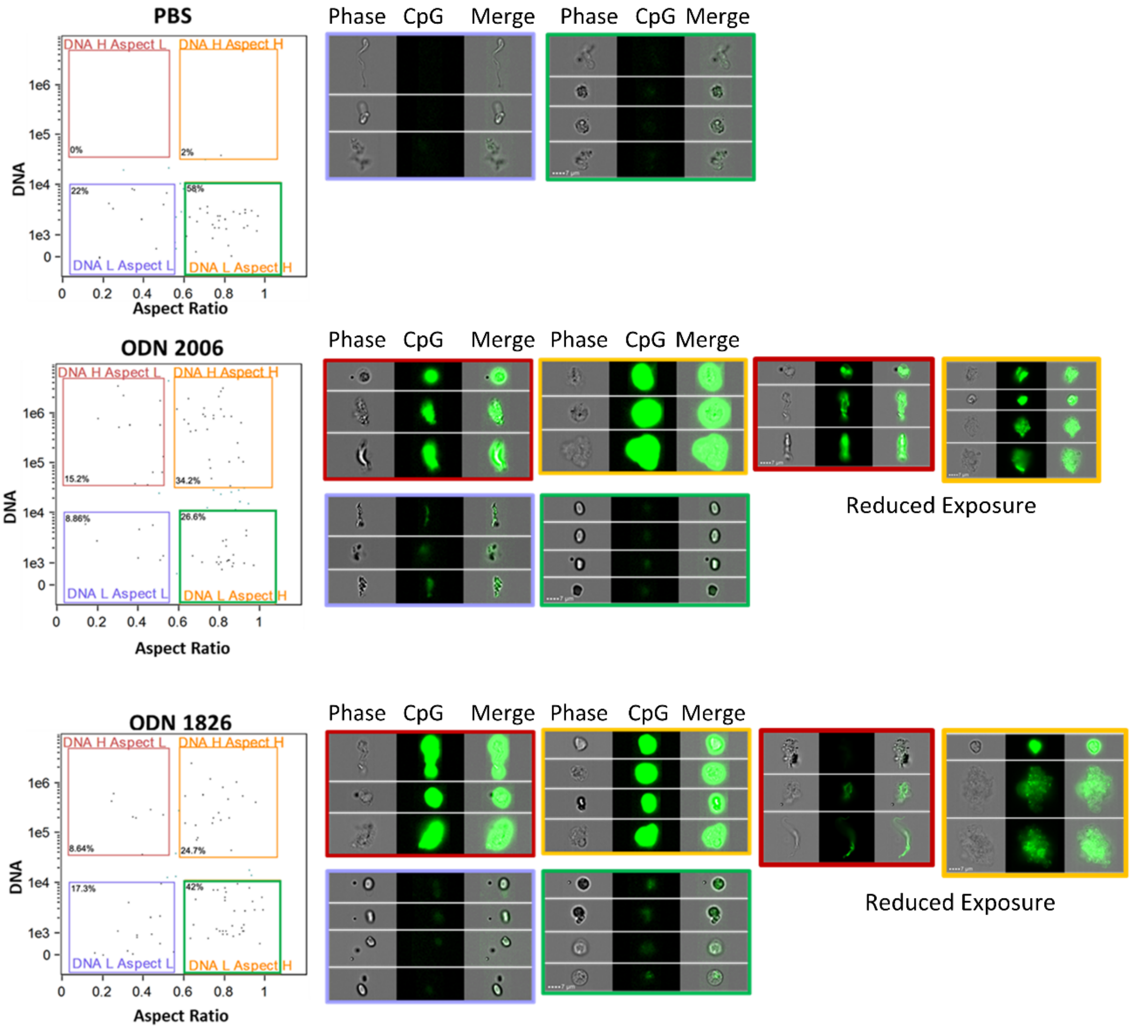


**Supplemental Figure 3. Imaging Flow Cytometry of CpG-treated Bat RBCs.** a. PBS treated RBCs b. RBCs treated with ODN 1826 (b) and ODN 2006 (c) reveal four distinct populations of cells and acquisition of DNA by RBCs. Reduced intensity images are provided for the DNA positive cells, right most panels.

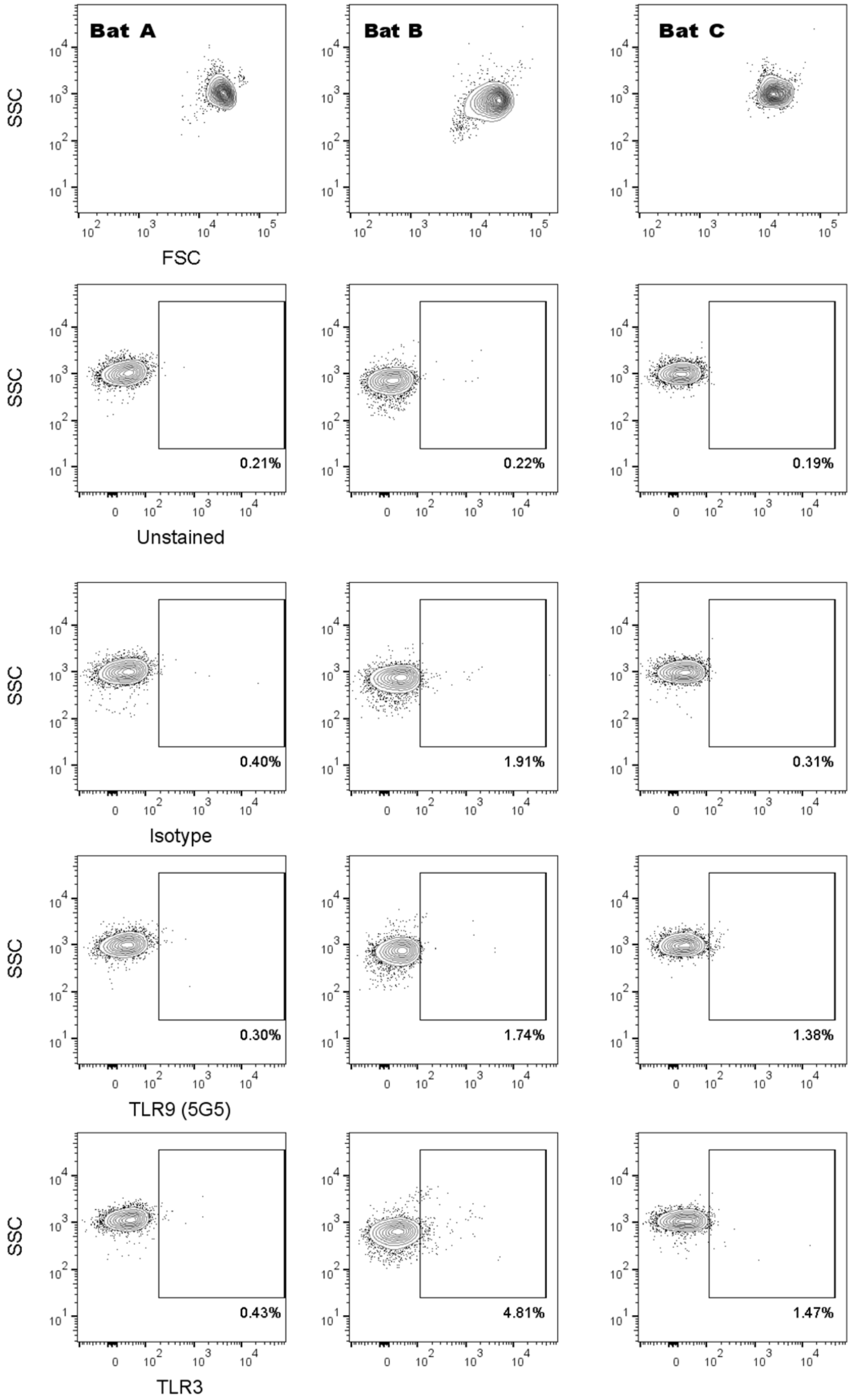


**Supplemental Figure 4. Surface staining for TLRs on Bat RBCs.** Flow cytometry for TLR 3 and 9 on intact, non-permeabilized RBCs from three representative bats.
